# ORF3c is expressed in SARS-CoV-2 infected cells and suppresses immune activation by inhibiting innate sensing

**DOI:** 10.1101/2023.02.27.530232

**Authors:** Martin Müller, Alexandra Herrmann, Shigeru Fujita, Keiya Uriu, Carolin Kruth, Adam Strange, Jan E. Kolberg, Markus Schneider, Jumpei Ito, Armin Ensser, The Genotype to Phenotype Japan (G2P-Japan) Consortium, Kei Sato, Daniel Sauter

**Affiliations:** Institute for Medical Virology and Epidemiology of Viral Diseases, University Hospital Tübingen, 72076 Tübingen, Germany; Institute for Clinical and Molecular Virology, University Hospital, Friedrich-Alexander-Universitat Erlangen-Nürnberg, 9l054 Erlangen, Germany; Division of Systems Virology, Department of Microbiology and Immunology, The Institute of Medical Science, The University of Tokyo, Minato-ku, Tokyo, Japan; Graduate School of Medicine, The University of Tokyo, Minato-ku, Tokyo, Japan; International Research Center for Infectious Diseases, The Institute of Medical Science, The University of Tokyo, Minato-ku, Tokyo, Japan; International Vaccine Design Center, The Institute of Medical Science, The University of Tokyo, Minato-ku, Tokyo, Japan; Graduate School of Frontier Sciences, The University of Tokyo, Kashiwa, Chiba, Japan; CREST, Japan Science and Technology Agency, Kawaguchi, Saitama, Japan

## Abstract

SARS-CoV-2 proteins are translated from subgenomic RNAs (sgRNAs). While most of these sgRNAs are monocistronic, some viral mRNAs encode more than one protein. For example, the *ORF3a* sgRNA also encodes ORF3c, an enigmatic 41-amino acid peptide. Here, we show that ORF3c is expressed in SARS-CoV-2 infected cells and suppresses RIG-I- and MDA5-mediated immune activation and IFN-β induction. Mechanistic analyses revealed that ORF3c interacts with the signaling adaptor MAVS, induces its C-terminal cleavage and inhibits the interaction of RIG-I with MAVS. The immunosuppressive activity of ORF3c is conserved among members of the subgenus sarbecovirus, including SARS-CoV and coronaviruses isolated from bats. Notably, however, the SARS-CoV-2 delta and kappa variants harbor premature stop codons in ORF3c demonstrating that this reading frame is not essential for efficient viral replication *in vivo* and likely compensated by other viral proteins. In agreement with this, disruption of ORF3c did not significantly affect SARS-CoV-2 replication in CaCo-2 or CaLu-3 cells. In summary, we here identify ORF3c as an immune evasion factor of SARS-CoV-2 that suppresses innate sensing in infected cells.

## INTRODUCTION

Since the emergence of the COVID-19 pandemic, all canonical proteins of SARS-CoV-2 have been extensively characterized for their expression, structure and function. In addition to its prototypical genes, however, SARS-CoV-2 harbors several smaller open reading frames (ORFs) that frequently overlap with other ORFs and may also contribute to efficient viral replication. For example, the ORF3b peptide encoded by *ORF3a* subgenomic RNA (sgRNA) was shown to suppress the induction of type I interferon (IFN) [1]. Intriguingly, naturally occurring variants of ORF3b differ in their immunosuppressive activity and may be responsible for phenotypic differences between SARS-CoV and SARS-CoV-2 [1]. Moreover, several short upstream ORFs (uORFs) have been suggested to regulate translation of downstream genes such as *ORF7b* [2]. Thus, non-canonical ORFs of SARS-CoV-2 may also be important determinants of viral immune evasion, spread and/or pathogenicity.

Nevertheless, most of the cryptic ORFs of SARS-CoV-2 remain poorly characterized, and several open questions remain: Do they encode proteins or are they merely a result of selection pressures acting on overlapping reading frames? Do these ORFs exert any regulatory activity, e.g. by modulating translation of downstream ORFs via leaky scanning or ribosomal re-initiation? Do they code for functional proteins that contribute to efficient immune evasion and/or replication of SARS-CoV-2? Are the respective peptides or proteins immunogenic? One interesting cryptic open reading frame is *ORF3c*, located at nt 25457-25579 of the Wuhan-Hu-1 reference genome. This ORF was independently described by different groups and has received several alternative names: ORF3c [3, 4], ORF3h (for hypothetical) [5], 3a.iORF1 [2], and ORF3b [6]. Following the homology-based nomenclature proposed by Jungreis and colleagues [7], we will refer to this open reading frame as *ORF3c* hereafter. Like *ORF3b*, *ORF3c* is one of several open reading frames overlapping with *ORF3a*. *In silico* analyses suggested that the respective ORF3c protein may harbor a transmembrane domain [3] and act as a viroporin [5]. Nevertheless, its expression in infected cells, as well as its exact function and relevance for viral replication have remained unclear.

Here, we show that SARS-CoV-2 ORF3c encodes a stable 41-amino acid peptide that is expressed in virally infected cells and suppresses the induction of IFN-β expression. Mechanistic analyses revealed that it inhibits innate sensing induced by RIG-I and MDA5, and interacts with the downstream adaptor protein MAVS. Furthermore, it inhibits the interaction of RIG-I with MAVS and induces proteolytic cleavage of MAVS. In line with a relevant function *in vivo*, ORF3c orthologs from different sarbecoviruses share this immunosuppressive activity. However, we also identify SARS-CoV-2 lineages that spread efficiently in the human population despite premature stop codons in their *ORF3c* genes. Furthermore, disruption of *ORF3c* did not affect SARS-CoV-2 replication in CaCo-2 and CaLu-3 cells. Thus, our findings identify ORF3c as an immune evasion factor that inhibits innate sensing cascades, but is not essential for efficient viral replication.

## RESULTS

### SARS-CoV-2 *ORF3c* encodes a peptide suppressing IFN-β promoter activation

The *ORF3a* gene of the SARS-CoV-2 reference genome Wuhan-Hu-1 overlaps with several shorter open reading frames that have the potential to encode for peptides of at least l0 amino acids in length (Fig 1A). Translation initiation downstream of the start codon of *ORF3a* may be enabled via non-canonical translation mechanisms, such as leaky scanning, ribosomal shunting and/or re-initiation [8]. In line with this, the start codon of ORF3c is part of a strong Kozak sequence, and *in silico* analyses predict ORF3c expression from the ORF3a sgRNA via leaky scanning [9] (Fig 1B). Indeed, ribosome profiling and HLA-ü immunopeptidome studies suggested that *ORF3c* is translated in SARS-CoV-2-infected cells [2, 10]. The same study also found evidence for translation of *ORF3d-2* [2], which is in agreement with the detection of ORF3d-specific antibodies in sera from previously SARS-CoV-2-infected individuals [11]. In contrast, we found no evidence for antibodies against ORF3c in SARS-CoV-2 convalescent sera (Fig Sl)

**Figure 1:**
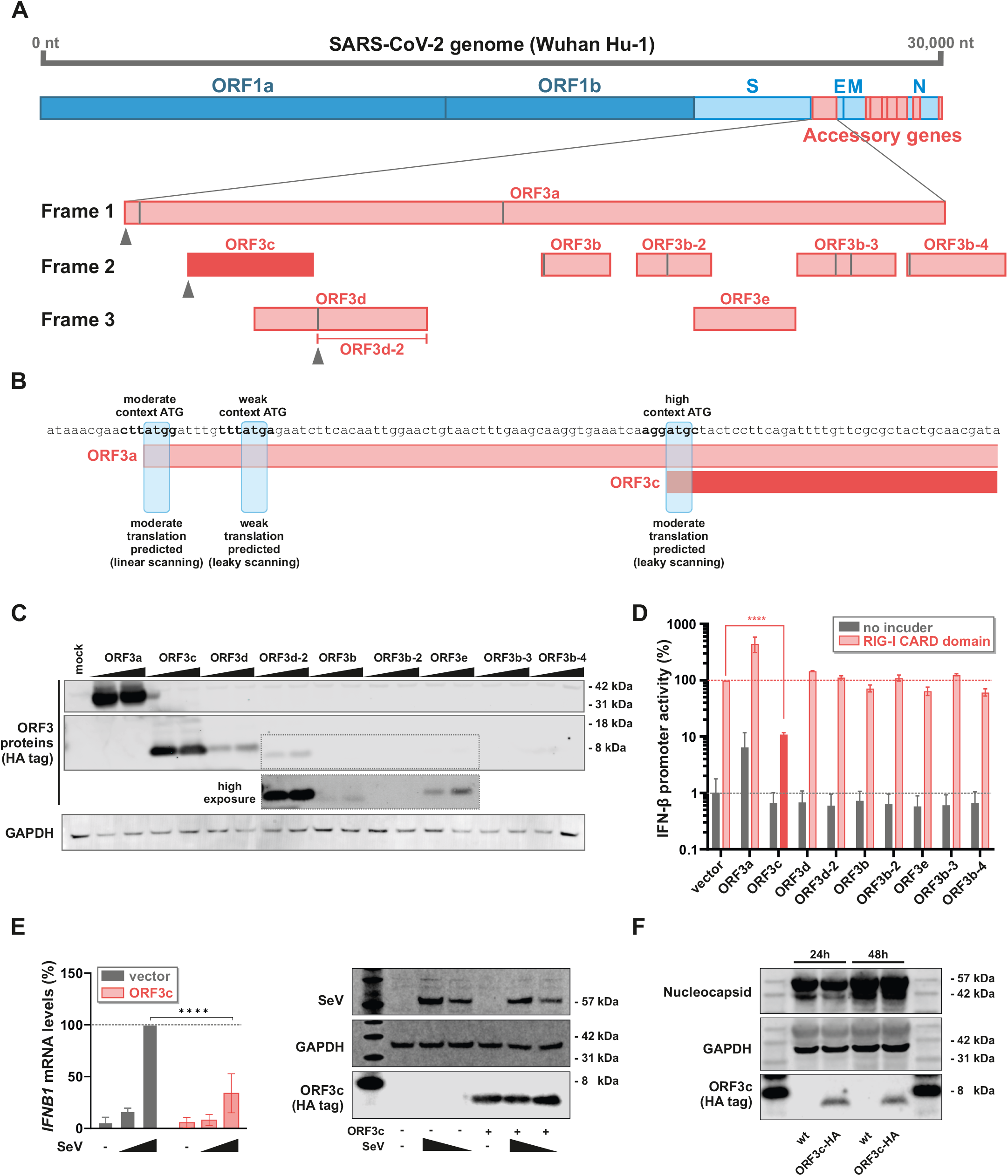
Open reading frames in the *ORF3* gene locus and inhibition of IFN-β induction by SARS-CoV-2 ORF3c. **A** Genome organization of SARS-CoV-2 is illustrated on top; overlapping ORFs in the ORF3 locus are shown at the bottom. Vertical grey lines indicate internal ATG codons. Experimentally confirmed translation initiation sites [2] are highlighted by grey triangles. **B** Kozak sequences of the *ORF3c* initiation codon and upstream ATG codons in *ORF3a* are shown. ATG context was determined using TIS predictor (https://www.tispredictor.com/) [9]. **C** Western blot analysis of HEK293T cells transfected with two different concentrations of expression plasmids for the indicated ORF3 proteins and peptides. After 24h ORF3a to ORF3e were detected via a C-terminal HA tag. GAPDH served as loading control. **D** HEK293T cells were co-transfected with the indicated ORF3 expression plasmids, a reporter plasmid expressing firefly luciferase under the control of the *IFNBl* promoter and a construct expressing *Gaussia* luciferase under the control of a minimal promoter. To induce immune signaling, half of the samples were additionally co-transfected with an expression plasmid for the CARD domain of RIG-I. One day post transfection, firefly luciferase activity was determined and normalized to *Gaussia* luciferase activity. **E** HEK293T cells were transfected with an expression plasmid for SARS-CoV-2 ORF3c or an empty vector control. 24 hours post transfection, cells were infected with increasing amounts of Sendai virus (SeV) for an additional 8 hours. Cells were lysed to perform either RNA extraction and subsequent qPCR for IFN-β (left panel) or Western blot analysis (right panel). **F** CaCo-2 cells were infected with SARS-CoV-2 wt or SARS-CoV-2 encoding HA-tagged ORF3c at an MOI of 2. 24 and 48 hours post infection, cells were harvested for Western blot analysis. ORF3c expression was detected via the HA tag. SARS-CoV-2 Nucleocapsid and GAPDH served as controls. Data information: In (D–E), data are presented as mean (±SD) of three independent experiments. Multiple comparison within individual reporter assays (D) were determined by one-way ANOVA with Dunnett’s test * p ≤ 0.05; ** p ≤ 0.01; *** p ≤ % 0.001 and **** p ≤ 0.000l. Multiple comparison between groups (E) were determined by two-way ANOVA with Sidak’s multiple comparison test; * p ≤ 0.05; ** p ≤ 0.01; *** p ≤ % 0.001 and **** p ≤ 0.0001.

To characterize the stability and potential activity of cryptic ORF3 peptides, we generated expression vectors for the individual peptides harboring a C-terminal HA-tag (without codon-optimization). Apart from the *ORF3a* construct, *ORF3c* and *ORF3d/ORF3d-2* code for stable proteins that are readily detectable in transfected cells (Fig 1C). The remaining peptides were not (ORF3b-2, ORF3b-3, ORF3b-4) or only poorly (ORF3b, ORF3e) detectable.

Since several accessory proteins of SARS-CoV-2 (e.g., ORF3b, ORF6) have been shown to suppress the induction of interferons [1, l2, l3], we hypothesized that some of the cryptic ORF3 peptides may exert similar immune-modulatory activities. Indeed, a luciferase reporter assay revealed that ORF3c significantly suppresses the activation of the IFN-β promoter in response to a constitutively active mutant of the pattern recognition receptor RIG-I (Fig 1D). Notably, ORF3c was also more active than the previously described IFN antagonist ORF3b, which suppressed IFN-β promoter activity only at higher concentrations or upon codon-optimization [1]. To test whether ORF3c is able to suppress immune activation upon viral infection, we monitored endogenous *IFNBl* expression upon infection with Sendai virus (SeV), a potent inducer of RIG-I-mediated type I IFN expression [14]. As expected, SeV induced *IFNBl* expression in a dose-dependent manner (Fig 1E). However, *IFNBl* mRNA levels were reduced by about 70% in the presence of SARS-CoV-2 ORF3c.

To definitely prove that ORF3c is produced in SARS-CoV-2-infected cells, we monitored its expression at different time points post infection via Western blot. Due to the lack of ORF3c-specific antibodies, we used circular polymerase extension reaction (CPER) [15] to generate a SARS-CoV-2 variant expressing C-terminally HA-tagged ORF3c. Although addition of the HA tag also resulted in the insertion of ten additional amino acids within ORF3a, the rescued virus grew to high titers, similar to the wt virus (not shown). Most importantly, ORF3c-HA was readily detectable in CaCo-2 cells, 24 h and 48 h post infection (Fig 1F). To our knowledge this is the first direct detection of ORF3c in SARS-CoV-2 infected cells. Together, these findings show that SARS-CoV-2 *ORF3c* encodes a stable peptide that suppresses the production of IFN-β upon viral sensing.

### ORF3c targets MAVS and suppresses both RIG-I- and MDAS-mediated immune activation

To elucidate the mechanisms underlying the inhibitory activity of ORF3, we analyzed different steps of the innate RNA sensing cascade culminating in the induction of IFN-β expression (Fig 2A). The *IFNB1* promoter harbors binding sites for both IRF3 and NF-κB. As expected, disruption of the NF-κB binding site reduced responsiveness to RIG-I-mediated activation (Fig 2B). However, ORF3c still dose-dependently reduced promoter activation (Fig 2B), demonstrating that ORF3c does not selectively target NF-κB activation. Next, we activated the sensing cascade at different steps via over-expressing MDA5, MAVS or a constitutively active mutant of IRF3. While ORF3c suppressed MDA5-mediated immune activation (Fig 2C, left panel), it failed to efficiently suppress *IFNB1* promoter activation in response to MAVS or IRF3 over-expression (Fig 2C, middle and right panels). Together, these findings suggest that ORF3c targets immune activation upstream or at the level of the signaling adaptor MAVS. In line with this, SARS-CoV-2 ORF3c weakly co-immunoprecipitated with MAVS, while we found no evidence for an interaction with RIG-I, MDA5 or TBK1 (Fig 2D). The modest co-immunoprecipitation of ORF3c was also observed when a MAVS mutant lacking its CARD domain was used for pull-down (Fig 2E), demonstrating that this domain is dispensable for the interaction. We hypothesized that the interaction of ORF3c with MAVS may also affect the binding of RIG-I to MAVS. Indeed, an established bimolecular fluorophore complementation assay [l6] showed evidence for reduced RIG-I:MAVS interaction in the presence of SARS-CoV-2 ORF3c (Fig 1F). Intriguingly, we also noted that increasing amounts of ORF3c resulted in the emergence of a C-terminal MAVS fragment of about 9 kDa (Fig 2G). Together, these findings suggest that ORF3c suppresses IFN-β expression by binding to MAVS, preventing its interaction with RIG-I, and/or inducing its C-terminal cleavage.

**Figure 2.**
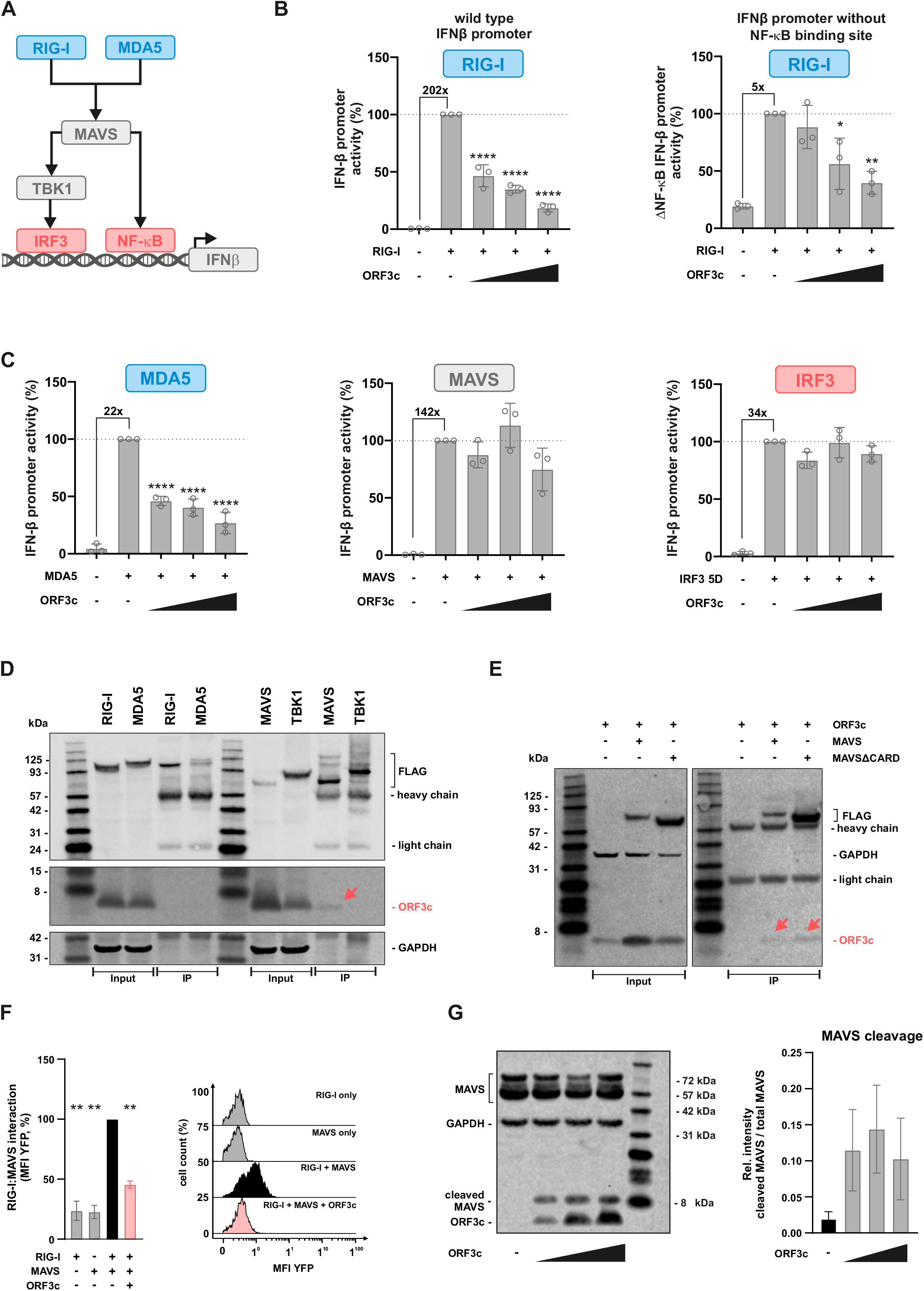
ORF3c interacts with MAVS and inhibits IFN-β induction independently of the pattern recognition receptor. **A** Cartoon illustrating IRF3- and NF-κB-mediated activation of the *IFNBl* promoter upon RIG-I- or MDA5-mediated sensing. **B** HEK293T cells were co-transfected with increasing amounts of an expression plasmid for SARS-CoV-2 ORF3c, a construct expressing *Gaussia* luciferase under the control of a minimal promoter and a reporter plasmid expressing firefly luciferase under the control of the *IFNBl* promoter (left panel) or a mutant thereof lacking the NF-κB binding site (right panel). Immune signaling was induced by co-transfecting an expression plasmid for the CARD domain of RIG-I. One day post transfection, firefly luciferase activity was determined and normalized to *Gaussia* luciferase activity. **C** HEK293T cells were transfected and analyzed essentially as described in (B). Immune signaling was introduced by co-transfecting expression plasmids for MDA5 (left panel), MAVS (central panel) or a constitutively active mutant of IRF3 (right panel). **D, E** HEK293T cells were co-transfected with expression plasmids for (D) Flag-tagged RIG-I, MDA5, MAVS, TBK1, (E) MAVS or a mutant thereof lacking its CARD domain (MAVSΔCARD) and an expression plasmid for HA-tagged SARS-CoV-2 ORF3c. One day post transfection, cells were lysed. Cell lysates were analyzed by Western blotting, either directly (“input”) or upon pull-down using a Flag-specific antibody (“IP”). **F** HEK293T cells were transfected with plasmids expressing BFP, ORF3c, the C-terminal part of YFP fused to RIG-I and/or the N-terminal part of YFP fused to MAVS. 24 h later, cells were fixed, and YFP fluorescence was detected by flow cytometry as a reporter for MAVS-RIG-I interaction. Exemplary flow cytometry data are shown on the right. **G** HEK293T cells were transfected with increasing amounts of an expression plasmid for SARS-CoV-2 ORF3c. One day post transfection, cells were lysed for Western blotting. ORF3c was detected via an HA tag, and MAVS was detected with antiserum specific for the C-terminal part of MAVS. GAPDH served as loading control. MAVS bands were quantified and the ratio of the 9 kDa fragment to total MAVS was calculated. One exemplary blot is shown on the left. Data information: In (B, C, F), data are presented as mean (±SD) of three independent experiments. In (G), data are presented as mean (±SEM) of three independent experiments. Multiple comparison within individual reporter assays were determined by one-way ANOVA with Dunnett’s test * p ≤ 0.05; ** p ≤ 0.01; *** p ≤ % 0.001 and **** p ≤ 0.0001.

### The immunosuppressive activity of ORF3c is conserved among sarbecoviruses

*In silico* analyses of the ORF3 locus revealed that essentially all sarbecoviruses harbor an *ORF3c* gene encoding a 40- or 41-amino-acid peptide [3]. In contrast, the remaining *ORF3* genes are only poorly conserved or vary substantially in their length (Fig 3A). The absence of *ORF3c* from other subgenera of betacoronaviruses suggests that this open reading frame emerged after the divergence of sarbeco- and hibecoviruses. To test whether the immunosuppressive activity of ORF3c is also conserved, we characterized several orthologs representing human and bat isolates of the SARS-CoV- and SARS-CoV-2-like clusters (Fig 3B). Titration experiments revealed that all ORF3c peptides tested significantly suppress IFN-β promoter activation (Fig 3C). However, ORF3c of SARS-CoV-2 Wuhan Hu-1 and a closely related bat coronavirus (ZXC21) suppressed IFN-β promoter activation more efficiently than ORF3c of SARS-CoV Tor2 and SARS-CoV-2 BANAL-20-50.

**Figure 3.**
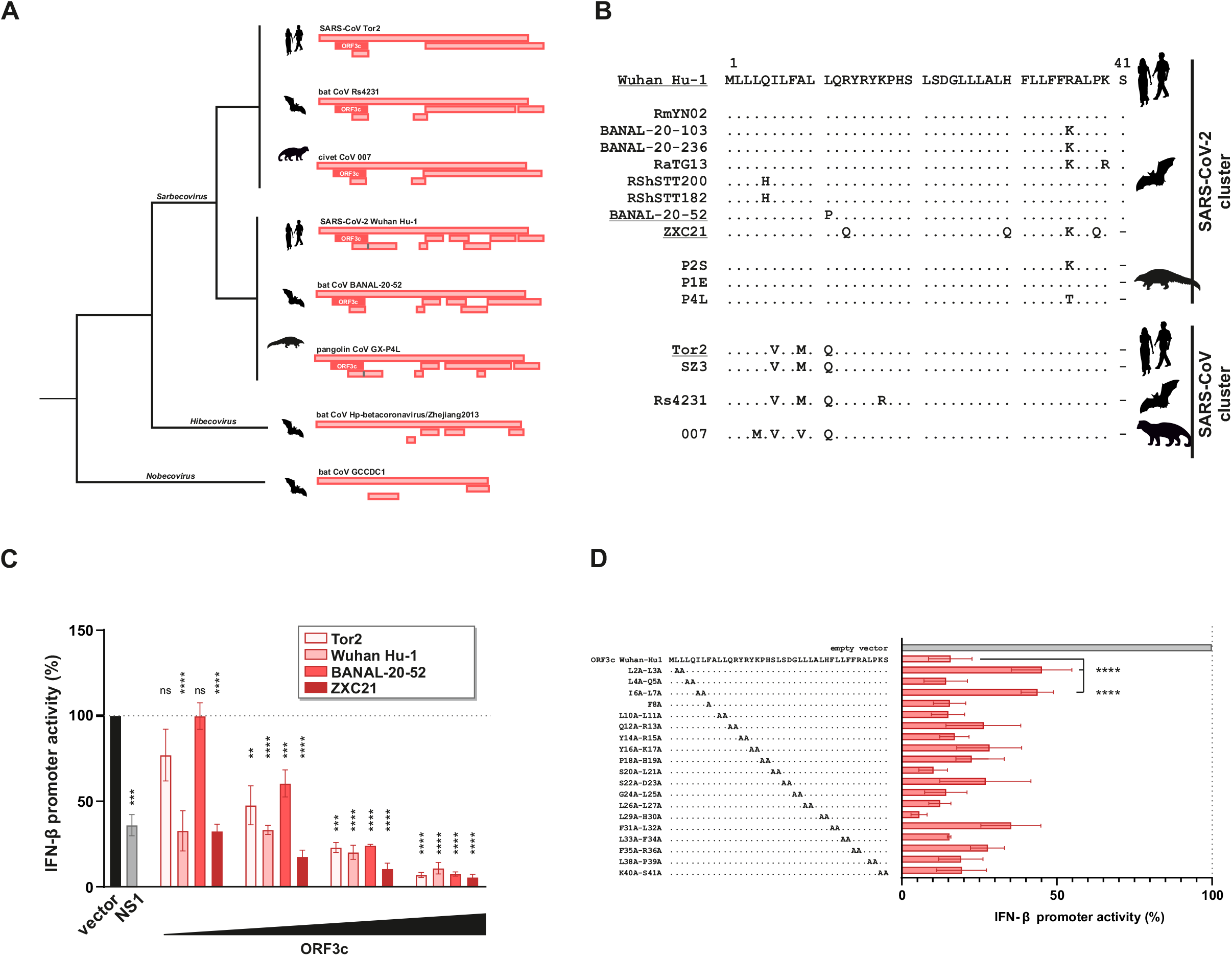
Conservation of ORF3c and its immunosuppressive activity in sarbecoviruses. **A** Simplified cartoon illustrating the *ORF3* locus of randomly selected members of the *Sarbeco*-, *Hibeco*- and *Nobecovirus* genera. Open reading frames with a length of at least 30 nucleotides are indicated as rectangles. ORF3c is highlighted in dark red. **B** Alignment of ORF3c amino acid sequences of the indicated viral isolates. Members of the SARS-CoV-2 cluster are shown on top, members of the SARS-CoV cluster at the bottom. For the underlined ORF3c sequences, expression plasmids were generated and analyzed for their ability to inhibit *IFNBl* promoter activation in **C** HEK293T cells were co-transfected with increasing amounts of the indicated ORF3c expression plasmids, a reporter plasmid expressing firefly luciferase under the control of the *IFNBl* promoter and a construct expressing *Gaussia* luciferase under the control of a minimal promoter. An expression plasmid for Influenza A virus non-structural protein l (NSl) served as positive control. Immune signaling was induced by co-transfecting an expression plasmid for the CARD domain of RIG-I. One day post transfection, firefly luciferase activity was determined and normalized to *Gaussia* luciferase activity. **D** HEK293T cells were co-transfected with expression plasmids and analyzed essentially as described in (C). The respective alanine mutations are indicated in the alignment on the left. Data information: In (C, D), data are presented as mean (±SD) of three independent experiments. Multiple comparison between groups were determined by two-way ANOVA with Sidak’s multiple comparison test; * p ≤ 0.05; ** p ≤ 0.01; *** p ≤ % 0.001 and **** p ≤ 0.0001.

To further map residues in ORF3c involved in its immunosuppressive activity, we performed an alanine scan using the Wuhan Hu-1 orthologue (Fig 3D). While all ORF3c mutants tested still reduced IFN-β promoter activity, the double mutants L2A/L3A and I6A/L7A significantly reduced the immunosuppressive effect of SARS-CoV-2 ORF3c. In summary, these results identify residues in the N-terminus of ORF3c that contribute to its inhibitory activity and demonstrate that ORF3c orthologs from different sarbecoviruses suppress the induction of IFN-β, albeit to different degrees.

### A natural R36I polymorphism does not affect the immunosuppressive activity of ORF3c

Since the emergence of SARS-CoV-2 in 2019, several mutations have occurred throughout the viral genome. One notable mutation is G25563T (Fig 4A), which is found in the beta, eta, iota and mu variants and has been associated with increased transmission fitness [17]. Although G25563T simultaneously introduces non-synonymous mutations in ORF3a (Q57H), ORF3c (R36I) and ORF3d (E3l*) (Fig 4A), previous studies focused only on possible phenotypic consequences of the Q57H change in ORF3a. Notably, however, the R36I mutation in ORF3c is predicted to result in a conformational change (Fig 4B) and a transmembrane domain in the C-terminal half of ORF3c (Fig 4C). We therefore analyzed whether the R36I change may affect the subcellular localization and/or immune-suppressive activity of SARS-CoV-2 ORF3c. Immunofluorescence microscopy revealed that ORF3c of the Wuhan-Hu-1 reference strain and a R36I mutant thereof are similarly distributed throughout the cytoplasm (Fig 4D). Moreover, both ORF3c variants dose-dependently suppressed RIG-I-mediated IFN-β promoter activation to a similar extent (Fig 4E). Thus, the G25563T polymorphism of the SARS-CoV-2 beta, eta, iota and mu variants does not seem to alter the IFN-suppressive activity of ORF3c.

**Figure 4.**
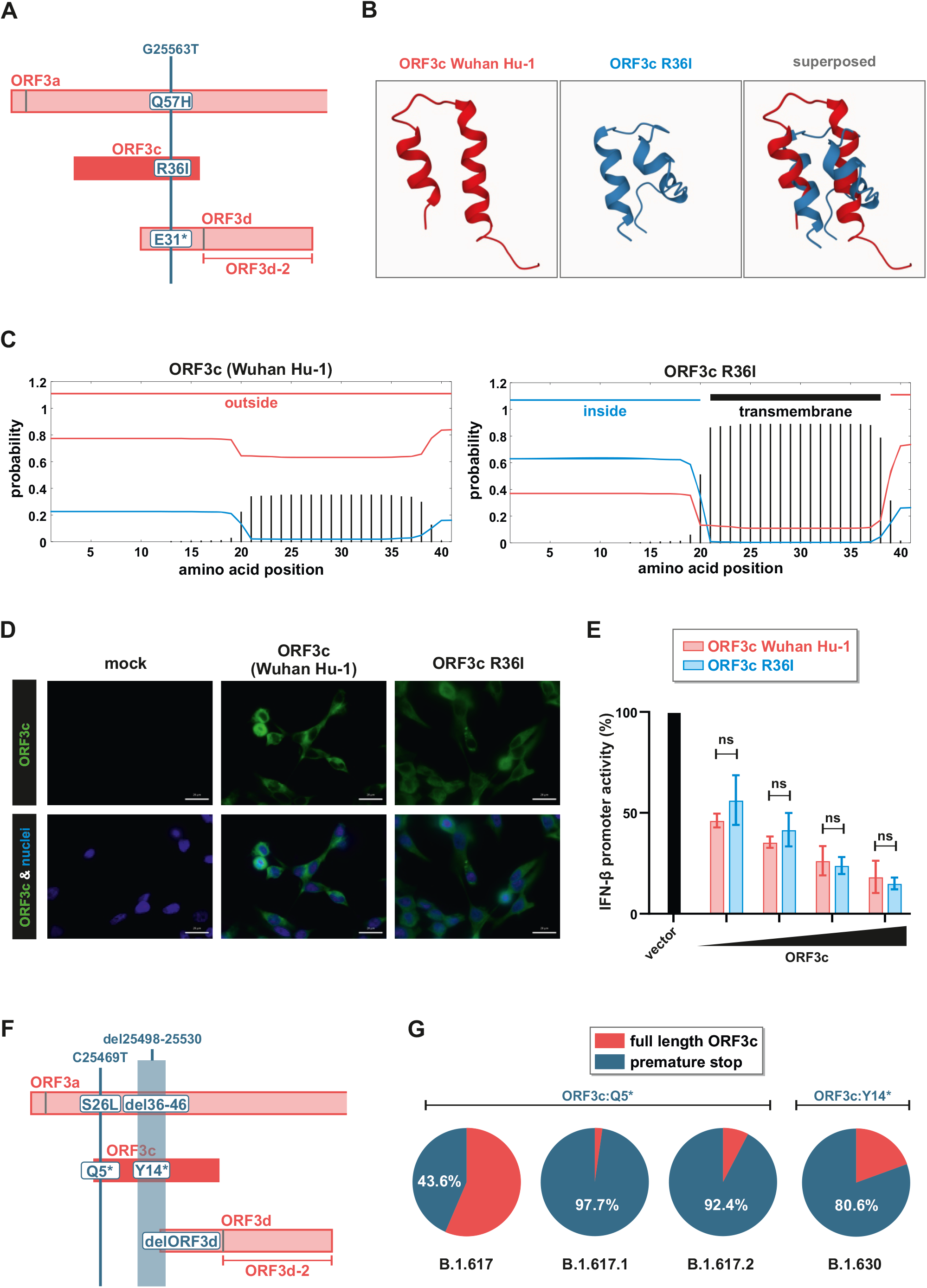
Characterization of naturally occurring variants of SARS-CoV-2 ORF3c. **A** Cartoon illustrating non-synonymous changes in ORF3a, c and d as a result of the naturally occurring polymorphisms G25563T. **B** Secondary structure of Wuhan Hu-l ORF3c (red) and the respective R36I variant thereof (blue) as predicted using PEP-FOLD 3 [32]. **C** The presence of transmembrane domains in Wuhan Hu-l ORF3c (left panel) and the respective R36I variant thereof (right panel) was predicted using TMHMM - 2.0 [33]. **D** HEK293T cells were transfected with expression plasmids for Wuhan Hu-l ORF3c or ORF3c R36I. One day post transfection, cells were stained for ORF3c (anti-HA, green) and nuclei (DAPI, blue) (scale bar = 20 µm). **E** HEK293T cells were co-transfected with increasing amounts of the indicated ORF3c expression plasmids, a reporter plasmid expressing firefly luciferase under the control of the *IFNBl* promoter and a construct expressing *Gaussia* luciferase under the control of a minimal promoter. Immune signaling was induced by co-transfecting an expression plasmid for the CARD domain of RIG-I. One day post transfection, firefly luciferase activity was determined and normalized to *Gaussia* luciferase activity. **F** Mutations introducing premature stop codons in ORF3c that can be found in at least 20% of the samples of at least one PANGO (sub)lineage. **G** Frequency of the mutations shown in (F) in the PANGO (sub)lineages B.l.6l7.l (delta), B.l.6l7.2 (kappa), B.l.6l7 and B.l.630. Data information: In (E), data are presented as mean (±SD) of three independent experiments. Multiple comparison between groups were determined by two-way ANOVA with Sidak’s multiple comparison test; * p ≤ 0.05; ** p ≤ 0.01; *** p ≤ % 0.001 and **** p ≤ 0.0001.

### Some SARS-CoV-2 variants harbor premature stop codons in *ORF3c*

To better understand the relevance of an intact *ORF3c* gene for viral spread, we screened the GISAID SARS-CoV-2 sequence repository for PANGO (sub)lineages harboring a premature ORF3c stop codon in at least 20% of the isolates. We identified two mutations fulfilling these criteria: the first one (C25469T) introduces a premature stop codon (Q5*) in ORF3c and an S26L change in ORF3a (Fig 4F). It is present in about 44% of B.l.6l7 isolates and in almost all sequences of the B.l.6l7.l (delta) and B.l.6l7.2 (kappa) sublineages (Fig 4G). The second mutation (del25498-25530) represents an in-frame deletion in ORF3a and can be found in about 80% of all B.1.630 isolates. This deletion results in the loss of the initiation codon of ORF3d and a premature stop codon (Yl4*) in ORF3c. The presence of premature ORF3c stop codons in a substantial fraction of B.l.6l7 and B.1.630 lineages suggests that ORF3c is dispensable for efficient viral replication *in vivo* and/or may be compensated by changes elsewhere in the genome. Notably, however, the analysis of GISAID SARS-CoV-2 sequences also revealed that a subset (~3%) of B.l.6l7.2 viruses had acquired an additional point mutation that reverted the stop codon at position 5 to a tyrosine (*5Y), thereby reconstituting ORF3c.

### ORF3c is dispensable for efficient SARS-CoV-2 replication

To assess the importance of *ORF3c* for efficient viral replication, we generated SARS-CoV-2 Wuhan Hu-1 variants harboring inactivating mutations in this gene. Introducing premature stop codons into *ORF3c* is not possible without simultaneously introducing non-synonymous mutations in ORF3a and/or ORF3d. We therefore mutated the start codon of ORF3c to threonine (M1T), which resulted in a silent mutation in the overlapping *ORF3a* gene (D22D) (Fig 5A, left panel). After rescue and validating successful introduction of the mutation, we infected CaCo-2 and CaLu-3 cells with an MOI of 0.l. Quantification of SARS-CoV-2 RNA copies in the culture supernatants over a period of three days revealed that ORF3c MlT replicated as efficiently as wild type SARS-CoV-2 (Fig 5A, right panels). In a parallel experiment, we introduced a premature stop codon (Q5*) in ORF3c, mimicking the mutation that can naturally be found in the SARS-CoV-2 delta and kappa variants (Fig 4F, 5B, left panel). Since the respective nucleotide change introduces a S26L mutation in ORF3a, we simultaneously disrupted *ORF3a* by introducing a premature stop codon (R6*) downstream of a methionine and potential alternative start codon at position 5. While loss of ORF3a markedly reduced replicative fitness in CaCo-2 cells, the virus still replicated efficiently in CaLu-3 cells (Fig 5B, right panels). As observed for the ORF3c M1T mutant, introduction of ORF3c Q5* did not significantly affect the replicative fitness of the virus.

**Figure 5.**
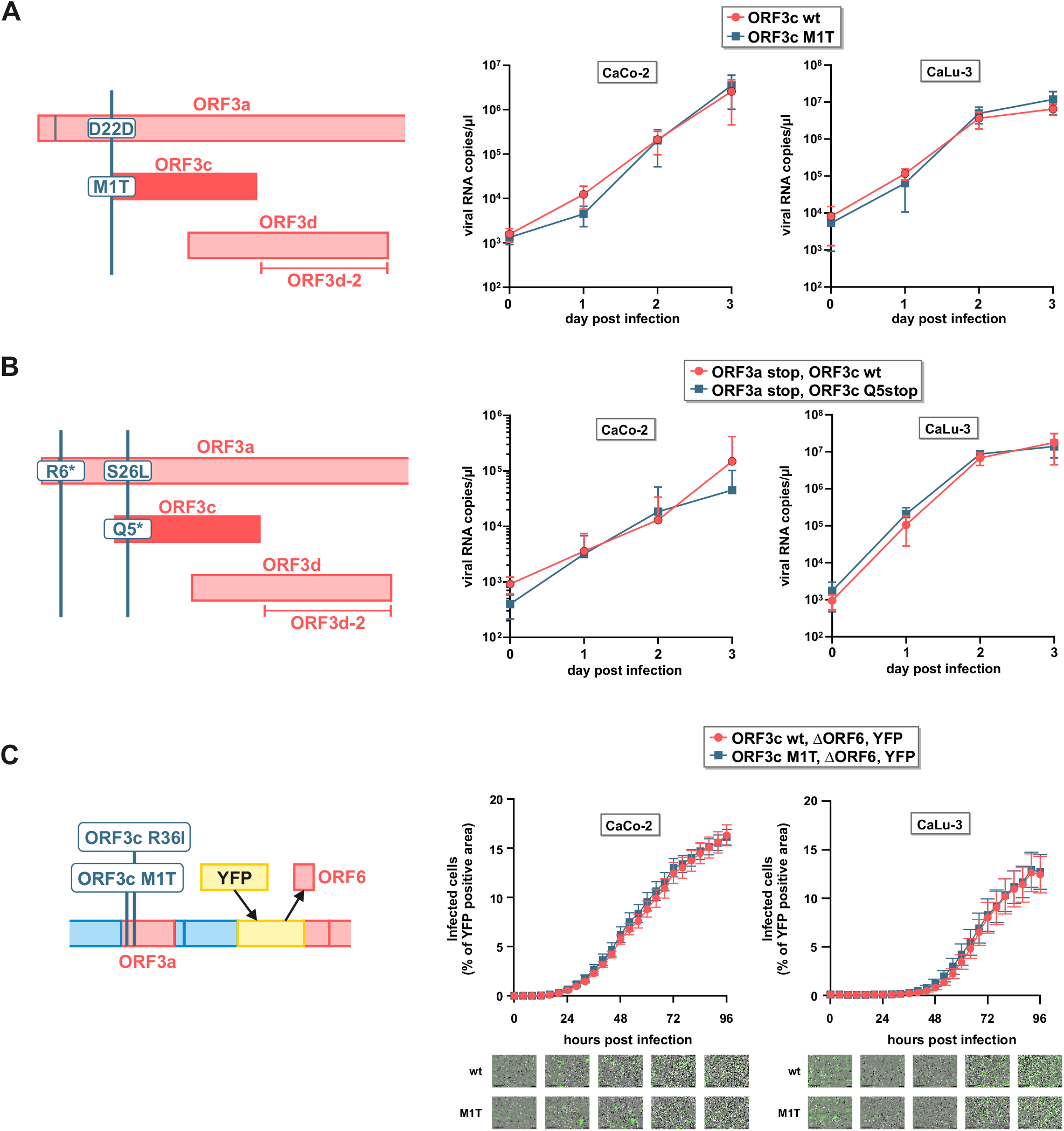
Disruption of ORF3c does not affect SARS-CoV-2 replication in CaCo-2 or CaLu-3 cells. **A** CPER was used to disrupt the start codon of ORF3c (MlT) in SARS-CoV-2 without affecting the amino acid sequence of ORF3a (left panel). CaCo-2 and CaLu-3 cells were infected with ORF3c wild type (red) or ORF3c-mutated (blue) SARS-CoV-2 at an MOI of 0.l. Viral replication was monitored over 72 hours by determining viral RNA copies in the culture supernatants (right panels). **B** CPER was used to introduce a premature stop codon in ORF3c (Q5stop). To avoid any bias by simultaneously changing the protein sequence of ORF3a (S26L), a premature stop codon was also inserted in ORF3a (R6*) (left panel). Viral replication (right panels) was monitored in CaCo-2 and CaLu-3 cells as described in (A). **C** A SARS-CoV-2 BAC clone harboring a disrupted ORF3c (MlT) and expressing YFP instead of ORF6 was generated (left panel). CaCo-2 and CaLu-3 cells were infected with SARS-CoV-2 ΔORF6-YFP (red) or SARS-CoV-2 ΔORF6-YFP ΔORF3c (blue) at an MOI of 0.l. Cells were placed in a live cell imaging device, and the area of YFP positive cells over the total area of cells was quantified every 4 hours for 96 hours. Images below graphs show continuous virus spread (green) over the indicated time points from one randomly chosen well. Data information: In (A-C, right panel), data are presented as mean (A, B ±SD and C ±SEM) of three independent experiments. Multiple comparison between groups were determined by two-way ANOVA with Sidak’s multiple comparison test; * p ≤ 0.05; ** p ≤ 0.01; *** p ≤ % 0.001 and **** p ≤ 0.0001.

Since several (accessory) proteins of SARS-CoV-2 have been shown to suppress the induction of IFN-β and/or IFN-β-mediated immune activation [l8, l9], we hypothesized that the immunosuppressive effect of ORF3c may be masked by other viral factors. One potent suppressor of IRF3-mediated IFN-β induction is ORF6 [12, 20]. We therefore generated a BAC clone of SARS-CoV-2, in which *ORF6* was replaced by a *YFP* reporter gene, and introduced the ORF3c M1T mutation described above (Fig 5C, left panel). This clone is based on the B.l variant, and therefore additionally harbors the ORF3c R36I mutation described above. Viral spread was monitored by live-cell imaging and quantification of YFP-expressing (i.e. infected) cells. Again, disruption of *ORF3c* did not result in impaired viral spread, and the ORF3c MlT virus replicated as efficiently as its ORF3c wild type counterpart (Fig 5C, right panels). Thus, while ORF3c and its immunosuppressive activity are conserved among sarbecoviruses, the respective gene is dispensable for viral replication in CaCo-2 and CaLu-3 cells.

## DISCUSSION

Several lines of evidence suggested that the *ORF3c* gene of SARS-CoV-2 codes for a peptide that plays a role in viral replication: (1) the respective open reading frame is conserved in different sarbecoviruses and shows synonymous site conservation [3, 4, 21] (Fig 3), (2) upstream ATGs do not show a strong initiation context and may allow ORF3c translation via leaky scanning [3] (Fig 1), (3) ribosomal profiling and the HLA-ü immunopeptidome suggested that ORF3c is translated in SARS-CoV-2 infected cells [2, 10], (4) ORF3c shows a high density of CD8+ T cell epitopes [21], (5) *in silico* analyses predict a conserved transmembrane domain in ORF3c and a potential role as viroporin [3, 5]. Still, the expression of ORF3c in infected cells, its exact function and its contribution to efficient viral replication have remained unclear. Here, we identify ORF3c as an immune evasion factor of SARS-CoV-2 and other sarbecoviruses that inhibits the induction of IFN-β upon activation of innate sensing cascades. The generation of a replication-competent SARS-CoV-2 variant encoding HA-tagged ORF3c allowed us to demonstrate the presence of a stable ORF3c peptide in virus infected cells. Using luciferase reporter assays, we show that ORF3c suppresses activation of the *IFNBl* promoter by the pattern recognition receptors (PRRs) and RNA sensors RIG-I and MDA5. Importantly, ORF3c also significantly reduced the expression of endogenous IFN-β in response to Sendai virus infection, a known inducer of RIG-I sensing [14]. Since ORF3c exerts its inhibitory effect independently of the receptor, it most likely targets a factor further downstream in the signaling cascade. In line with this, co-immunoprecipitation experiments revealed an interaction of ORF3c with the mitochondrial signaling adaptor MAVS. We found no evidence for an interaction with other components of the sensing cascade (i.e. RIG-I, MDA5, or TBK1). Notably, ORF3c failed to prevent *IFNB1* promoter activation if MAVS itself was used as an activator. This lack of inhibition is not merely the result of a saturation effect since MAVS induced the *IFNB1* promoter less efficiently than RIG-I (Fig 2). MAVS is targeted by proteins from different viruses. For example, Influenza A Virus PB1-F2 inhibits innate sensing by binding to MAVS [22] and decreasing the mitochondrial membrane potential [23]. Another example is the NS3/4A serine protease of Hepatitis C virus (HCV), which cleaves MAVS, thereby inhibiting downstream immune activation [24]. Similarly, SARS-CoV-2 ORFl0 was recently shown to induce the degradation of MAVS via mitophagy [25]. Further mechanistic analyses revealed that ORF3c induces the proteolytic processing of MAVS, resulting in a C-terminal 9 kDa fragment of this signaling protein (Fig 2G). Furthermore, ORF3c reduced the interaction of MAVS with RIG-I (Fig 2F). Thus, one possible mode of action is a competitive binding of ORF3c and RIG-I (and possible MDA5) to MAVS. In the presence of ORF3c, the CARD domain of MAVS may not be accessible and thus not be bound and activated by active RIG-I or MDA5. Notably, however, co-immunoprecipitation experiments showed that the CARD domain of MAVS is dispensable for an interaction of ORF3c with MAVS (Fig 2E). In line with a relevant role of ORF3c in viral replication, its immunosuppressive activity is conserved in orthologs of other sarbecovirus species, including the SARS-CoV reference virus Tor2. While all orthologs tested inhibited *IFNB1* promoter activation, those of SARS-CoV Tor2 and batCoV BANAL20-52 were less active than their counterparts from SARS-CoV-2 Wuhan-Hu-l and batCoV ZXC21. The reduced activity of BANAL-20-52 ORF3c can be ascribed to a single amino acid change (L11P) distinguishing it from Wuhan-Hu-1 ORF3c (Fig 3B). While Lll is largely conserved in the SARS-CoV-2 cluster, most of the viruses in the SARS-CoV cluster (including Tor2) harbor a glutamine at this position (Fig 3B) [3, 5]. Thus, polymorphisms at position ll can affect the inhibitory activity of ORF3c. In addition to this, our alanine scanning approach revealed that Leu2/Leu3 and Ile6/Leu7 are also contributing to the immunosuppressive effect of ORF3c (Fig 3D). Most of these residues are conserved among different sarbecoviruses. One notable exception is Ile6 (Fig 3B). Viruses from the SARS-CoV cluster harbor a Valine at this residue. These include SARS-CoV Tor2 ORF3c, which was less active than its SARS-CoV-2 counterpart.

ORF3c is not the only SARS-CoV-2 protein that interferes with RIG-I- and/or MDA-5-mediated immune activation. As already mentioned above, ORF10 suppresses innate sensing by targeting MAVS [25]. Moreover, ORF3b, nucleocapsid, ORF6 and ORF8 have all been shown to suppress IFN-β expression [1, 12, 20, 26, 27], highlighting the selection pressure exerted by this pathway. The convergent evolution of viral proteins exerting overlapping immune evasion activities may represent a backup mechanism that allows viral replication even if one of the IFN-β suppressing proteins is lost. In line with this, SARS-CoV-2 variants expressing a C-terminally truncated, inactive ORF6 protein have emerged several times during the pandemic and spread via human-to-human transmission [12]. Similarly, natural SARS-CoV-2 variants lacking an intact ORF3c gene still efficiently spread in the human population. In fact, more than 80% of the sequenced genomes of B.l.6l7.l, B.l.6l7.2 and B.l.630 harbor premature stop codons at positions 5 and 15, respectively (Fig 4). We hypothesized that the loss of ORF3c in these viruses may be compensated by the activity of ORF6. However, replication kinetics in CaCo-2 and CaLu-3 cells revealed that loss of ORF3c does not affect viral replication in the absence of ORF6 either (Fig 5). Intriguingly, IFN-β mRNA levels were not increased upon infection with the SARS-CoV-2 double mutant lacking ORF3c and ORF6 (data not shown). Thus, yet another viral inhibitor of IFN-β expression (e.g. ORF8 or N) may be able to rescue efficient viral replication in this case.

Notably, the emergence of premature stop codons in small reading frames such as *ORF3c* may also be tolerated or even be beneficial if they provide a fitness advantage by optimizing overlapping reading frames. For example, the ORF3c Q5* mutation is accompanied by an S26L change in ORF3a. However, experimental disruption of ORF3c without changing the amino acid sequence of ORF3a (Fig 5A) or upon deletion of ORF3a (Fig 5B) did not affect viral replication *in vitro* either.

One intriguing observation is the emergence of a nucleotide change in a subfraction (~3%) of B.l.6l7.2 viruses that reverts the stop codon at position 5 to a tyrosine (*5Y). Thus, it is tempting to speculate that the loss of ORF3c was initially just carried along with mutations elsewhere in the genome (e.g. in Spike) that conferred a major fitness advantage to the virus, before ORF3c expression was reverted by another point mutation.

In summary, our study identifies ORF3c as an immune evasion factor of SARS-CoV-2 and other sarbecoviruses. While an intact ORF3c gene is clearly dispensable for viral replication *in vitro* and *in vivo*, the conservation of this short open reading frame and the pseudo-reversion of premature stop codons suggests that it may still contribute to efficient viral replication *in vivo*. The emergence of future SARS-CoV-2 variants may help to fully decipher the role of this enigmatic ORF and its co-evolution with other viral genes.

## MATERIAL & METHODS

### Generation of expression plasmids

ORF3c genes were PCR-amplified using viral cDNA as a template and subsequently inserted into a pCG expression vector co-expressing GFP via an IRES using unique XbaI and MluI restriction sites. To facilitate protein detection, a C-terminal HA-tag (TACCCATACGATGTTCCAGATTACGCT) was added by extension PCR. To generate ORF3c-HA alanine mutants as well as ORF3c-R36I,-BANAL-20-52,-Tor2 and SL-CoVZXC2l, point mutations were introduced by site-directed mutagenesis using the wild type Wuhan-Hu-1 ORF3c-HA expression plasmid as template. pRen2-ORF3c was generated by conventional cloning, using the unique EcoRI and XhoI restriction sites in pRen2. IRF3 5D was PCR-amplified using pCAGGS Flag-IRF3 5D as a template and subsequently inserted into pEGFP-Cl via XhoI and EcoRI. All constructs were sequenced to verify their integrity. PCR primers are listed in EVl.

### Generation and recovery of a recombinant SARS-CoV-2 ORF3 mutant

Stop mutations within ORF3c were introduced into the bacmid pBSCoV2_d6-YFP harboring the SARS-CoV-2 backbone [28] using 2-step Red Recombination [29]. For this purpose, the KanS cassette was amplified from pEP-KanS with the following oligonucleotides:

1. mut3c-fwd: caattggaactgtaactttgaagcaaggtgaaatcaaggaCgctactccttcagattttgAGGATGACG ACGATAAGTAGGG
2. mut3c-rev: gtatcgttgcagtagcgcgaacaaaatctgaaggagtagcGtccttgatttcaccttgctCAACCAATTA ACCAATTCTGATTAG

Integrity of the obtained bacterial artificial chromosomes (BAC) and presence of desired stop mutations were confirmed by restriction digestion and next generation sequencing. Recombinant SARS-CoV-2 viruses expressing EYFP instead of the viral ORF6 protein and containing mutations within ORF3c were recovered by transfection of the BACs into HEK293T cells overexpressing viral N protein, ACE2 receptor, and T7 RNA polymerase as described previously [28]. The obtained reporter viruses were further passaged on CaCo-2 cells and viral titers were determined by endpoint titration (see TCID_50_).

### Generation of recombinant SARS-CoV-2 mutants by circular polymerase extension reaction

To generate recombinant SARS-CoV-2 by circular polymerase extension reaction (CPER) [l5] nine DNA fragments comprising parts of SARS-CoV-2 (WK-521, PANGO lineage A; GISAID ID: EPI ISL 408667) [30] were generated by PCR using PrimeSTAR GXL DNA polymerase (TAKARA, Cat# R050A). A linker fragment comprising hepatitis delta virus ribozyme, the bovine growth hormone poly A signal and the cytomegalovirus promoter was also prepared by PCR. The ten obtained DNA fragments were mixed and used for CPER. ORF3c mutations were inserted in fragment 9/10 by site-directed overlap extension PCR with the primers listed in EVl. To produce chimeric recombinant SARS-CoV-2, Tetracycline-inducible ACE2 and TMPRSS-expressing IFNARl-deficient HEK293 (HEK293-C34) cells were transfected with the CPER products using TransIT-LT1 (MirusBio, Cat#2300) according to the manufacturer’s protocol. One day post transfection, the culture medium was replaced with Dulbecco’s modified Eagle’s medium (high glucose) containing 2% FCS, 1% PS and doxycycline. At 7 d post transfection, the culture medium was harvested and centrifuged, and the supernatants were collected as the seed virus. To remove the CPER products (i.e., any SARS-CoV-2 DNA), 1 ml of the seed virus was treated with 2 μl TURBO DNase (Thermo Fisher Scientific, Cat# AM2238) and incubated at 37°C for 1 h. Complete removal of the CPER products (i.e., SARS-CoV-2-related DNA) from the seed virus was verified by PCR. To prepare virus stocks for infection, VeroE6/TMPRSS2 cells (5,000,000 cells in a T-75 flask) were infected with 20-50 µl of the seed virus. One-hour post infection, the culture medium was replaced with DMEM (low glucose) containing 2% FBS and 1% PS. Two to four days post infection, the culture medium was harvested and centrifuged, and the supernatants were collected. Viral titers were determined by TCID_50_. To verify the sequence of chimeric recombinant SARS-CoV-2, viral RNA was extracted from the virus stocks using the QIAamp viral RNA mini kit (Qiagen, Cat#74l36) and viral genomes were sequenced as described before [31].

### Tissue culture infectious dose (TCID_50_)

Viral titers were determined as the 50% tissue culture infectious dose. Briefly, one day before infection, VeroE6/TMPRSS2 cells (10,000 cells) were seeded into 96-well plates. Cells were inoculated with serially diluted virus stocks and incubated at 37°C. Four days later, cells were checked microscopically for cytopathic effects (CPE), and TCID_50_/ml was calculated using the Reed–Muench method.

### SARS-CoV-2 replication kinetics in Vero E6, Caco-2 and Calu-3 cells

One day before infection with CPER-derived SARS-CoV-2 clones, Caco-2 cells (l0,000 cells/well) or CaLu-3 cells (20,000 cells/well) were seeded into a 96-well plate. Cells were infected with SARS-CoV-2 at an MOI of 0.1 and incubated at 37°C. One hour later, the infected cells were washed and 180 μl of culture medium was added. The culture supernatants and cells were harvested at the indicated timepoints and used for RT–qPCR to quantify the viral RNA copy number. For replication kinetics of BAC-derived SARS-CoV-2 ΔORF6-YFP and SARS-CoV-2 ΔORF6-YFP ΔORF3c, Caco-2 cells (10,000/well) were seeded into a 96-well plate one day prior to infection. Cells were infected in triplicates at an MOI of 0.l & 0.0l for l hour at 37°C. After washing and addition of 100 µl fresh culture medium the plates were placed in an Incucyte plate reader and images were taken at the indicated time points for up to 96 hours. The ‘Basic Analysis Mode’ was applied to quantify virus growth as green area normalized to phase area. Supernatants and cells were harvested at the indicated time points to determine cytokine levels by Cytokine Array and RT-qPCR respectively.

### RT–qPCR

5 µl culture supernatant was mixed with 5 μL of 2 x RNA lysis buffer [2% Triton X-l00, 50 mM KCl, 100 mM Tris-HCl (pH 7.4), 40% glycerol, 0.8 U/μL recombinant RNase inhibitor and incubated at room temperature for 10 minutes. RNase-free water (90 μL) was added, and the diluted sample (2.5 μl) was used as the template for real-time RT-PCR performed according to the manufacturer’s protocol using the OneStep TB Green PrimeScript PLUS RT-PCR kit (TAKARA, Cat# RR096A) and the following primers: Forward *N*, 5′-AGC CTC TTC TCG TTC CTC ATC AC-3′; and Reverse *N*, 5′-CCG CCA TTG CCA GCC ATT C-3′. The viral RNA copy number was standardized using a home-made standard.

*IFNBl* and *GAPDH* RNA levels were determined in cell lysates collected from (1) SARS-CoV-2 ΔORF6-YFP and SARS-CoV-2 ΔORF6-YFP ΔORF3c infected CaCo-2 wild-type cells and (2) transfected HEK293T cells infected with Sendai virus (Cantell Strain) (Charles River, l0l00774) for 8 hours. Total RNA was isolated using the Viral RNA Mini Kit (Qiagen, Cat#74136) according to the manufacturer’s instructions. Genomic DNA was removed using the DNA-free kit (Thermo Fischer Scientific, Cat# AM1906) and subsequent cDNA synthesis was performed using the PrimeScript RT reagent Kit (TAKARA, Cat# RR037A), both according to the manufacturer’s instructions. qPCR was performed using the Luna Universal Probe qPCR Master Mix (NEB) together with primer probes for IFN-β (Thermo Fischer Scientific, Cat# 433ll82) and GAPDH (Thermo Fischer Scentific, Cat# 4448489). All reactions were run in duplicates and RNA levels were internally normalized to GAPDH.

### Transfection of HEK293T cells

For overexpression experiments, HEK293T cells were transfected using standard calcium phosphate transfection protocols. 6x l0^5^ cells were seeded in 6-well plates on the day before and medium was changed 6 hours after transfection.

### Bimolecular fluorescence complementation (BiFC) Assay

The BiFC assay was performed essentially as previously described [16]. In brief, HEK293T cells were transfected with a combination of expression plasmids encoding YFP amino acids l to l54 (yn) or 155 to the end (yc) conjugated to the N-terminus of RIG-I or MAVS, respectively. Cells were additionally transfected with an expression plasmid encoding different variants of ORF3c-HA or an empty vector control as well as an expression plasmid encoding BFP as transfection control (overall ratio: 2:2:1:0.5). After 24 h, cells were fixed using the Fix & Perm Kit l000 (Nordic-MUbio) according to the manufacturer’s instructions. ORF3c-HA was detected using a primary antibody against the HA-tag (Sigma-Aldrich, Cat# H3663) followed by staining with a secondary AF555 antibody (Invitrogen, Cat# A-21422). Samples were measured on a MACS Quant VYB (Miltenyi Biotec). Analysis was performed using FlowLogic V.8 (Inivai) and protein : protein interaction was determined as the MFI of YFP in BFP positive cells.

### Antibodies

The following primary antibodies were used, Rabbit polyclonal (IgG) anti-HA (l:l000, Sigma-Aldrich, Cat# SAB4300603), Mouse monoclonal anti-HA (l:l000, Sigma-Aldrich, Cat# H3663), Mouse monoclonal (IgG1) anti-Flag (l:l000, Sigma-Aldrich, Cat# Fl804), Rabbit polyclonal (IgG) anti-Flag (l:l000, Sigma-Aldrich, Cat# SAB430ll35), Rat polyclonal (IgG2a, κ) anti-human GAPDH (l:l000, BioLegend, Cat# 607902), Rabbit polyclonal anti-Sendai Virus (1:2000, MBL Life sience, Cat# PD029), Rabbit monoclonal (IgG) anti-RIG-I (l:l000, Cell Signaling, Cat# 3743), Rabbit monoclonal (IgG) anti-MDA5 (l:l000, Cell Signaling, Cat# 5321), Rabbit monoclonal (IgG) anti-MAVS (l:l000, Cell Signaling, Cat# 3993), Rabbit monoclonal (IgG) anti-TBKl (l:l000), Cell Signaling, Cat# 3504).

The following secondary antibodies were used for immunostaining, IRDye® 680RD Goat anti-Mouse IgG (H + L) (1:20000, LI-COR, Cat# 926-68070), IRDye® 680RD Goat anti-Rabbit IgG (H + L) (1:20000, LI-COR, Cat# 926-6807l), IRDye® 800CW Goat anti-Mouse IgG (H + L) (1:20000, LI-COR, Cat# 926-322l0), IRDye® 800CW Goat anti-Rabbit IgG (H + L) (l:20000, LI-COR, Cat# 926-322ll), IRDye® 800CW Goat anti-Streptavidin IgG (H + L) (l:20000, LI-COR, Cat# 926-32230), Alexa Fluor 488 goat anti rabbit IgG (1:250, Invitrogen, Cat# A-ll008), Alexa Fluor 488 goat anti mouse IgG (1:250, Invitrogen, Cat# A-ll00l), Alexa Fluor 555 goat anti mouse IgG (1:250, Invitrogen, Cat# A-21422), Goat anti-Rabbit IgG (H+L) Cross-Adsorbed Secondary Antibody, Alexa Fluor 555 (1:250, ThermoFisher, Cat# A-2l428).

### Co-immunoprecipitation

To investigate possible interactions between ORF3c and proteins of the Interferon signaling pathway, co-immunoprecipitation with subsequent analysis by Western blotting was performed. Briefly, HEK293T cells were seeded in 6-well plates and co-transfected with expression plasmids for HA-tagged ORF3c and Flag-tagged RIG-I, MDA5, MAVS or TBKl (ratio 4:1; 5µg/well). One day post transfection, cells were lysed in 300 µl Western blot lysis buffer and cleared by centrifugation (see “Western blotting”). 45 µl of the lysate was used for whole-cell lysate analysis and further prepared as described in “Western blotting”, while 255 µl of the lysate was used for co-immunoprecipitation. A pre-clearing step was performed to remove unspecifically binding compounds from the lysate. Pierce Protein A/G Magnetic beads (Thermo Fisher, Cat# 88802) were washed three times with 1 ml NP40 wash buffer (50 mM HEPES, 300 mM NaCl, 0.5% NP40, pH 7.4) and added to the lysate. After incubation for l h at 4°C, beads were removed from the lysate using a magnetic rack. To precipitate protein complexes, the lysate was incubated first with an anti-Flag antibody (1.5 µg/sample) for l hour followed by addition of 15 µl washed Protein A/G Magnetic Beads for one additional hour at 4°C. After incubation, the beads were washed three times in NP40 wash buffer before incubation with 80 µl 1 x Protein Sample Loading Buffer (LI-COR, Cat# 928-40004) at 95°C for 10 minutes to recover bound proteins. After addition of 1.75 ml β-mercaptoethanol, whole-cell lysates and precipitates were analyzed by western blotting.

### Western blotting

To determine expression of cellular and viral proteins, cells were washed in PBS, lysed in Western blot lysis buffer (150 mM NaCl, 50 mM HEPES, 5 mM EDTA, 0.1% NP40, 500 mM Na3VO4, 500 mM NaF, pH 7.5) and cleared by centrifugation at 20,800 x g for 20 min at 4°C. Lysates were mixed with Protein Sample Loading Buffer (LI-COR, Cat# 928-40004) supplemented with 10% β-mercaptoethanol and heated at 95°C for 5 min. Proteins were separated on NuPAGE 4%–12% Bis-Tris Gels (Thermo Fischer Scientific, Cat# NP0323BOX), blotted onto Immobilon-FL PVDF membranes (Merck Milipore, Cat# IPFL000l0) and stained using primary antibodies directed against HA-tag, Flag-tag, GAPDH, RIG-I, MDA5, MAVS, TBK1, IRF3 and Infrared Dye labeled secondary antibodies (LI-COR IRDye). Proteins were detected using a LI-COR Odyssey scanner and band intensities were quantified using LI-COR Image Studio Lite Version 5.2.

### Immunofluorescence microscopy

Confocal immunofluorescence microscopy was used to determine the subcellular localization of ORF3c and ORF3c R36I. Briefly, 150,000 HEK293T cells were seeded on 13 mm diameter glass coverslips coated with poly-L-lysine (Sigma-Aldrich, Cat# P6282-5mg) in 24-well plates. On the following day, cells were transfected with an expression plasmid for ORF3c, ORF3c R36I or an empty vector control (500 ng) using Lipofectamin2000 (Invitrogen, Cat# ll6680l9). One day post transfection, cells were fixed in 4% PFA for 20 min at RT, permeabilized in PBS 0.5% Triton X-l00 for l5 min at RT and blocked in 5% BSA/PBS supplemented with 0,l% Triton X-l00 for l5 min at RT. ORF3c was stained using a primary antibody against the HA-tag and secondary goat anti-mouse AF555. Nuclei were stained in parallel using 4′,6-Diamidine-2′-phenylindole (DAPI; Thermo Fischer Scientific, Cat# 62247). Coverslips were mounted on glass slides using Mowiol (CarlRoth, Cat# 07l3.l) mounting medium and confocal microscopy was performed using an LSM710 (Carl Zeiss).

### Firefly luciferase assay

HEK293T cells were seeded in 96 well plates at 3xl0^4^ cells /well. After 24 h, cells were transfected with a mix of expression vectors containing firefly luciferase reporter constructs for the IFN-β promoter or a mutant thereof lacking NF-κB biding sites (IFN-β ΔNF-κB) (reporter, 10 ng), a *Gaussia* luciferase expression plasmid (normalization control, 5 ng), expression plasmids for RIG-I-CARD, MDA5, MAVS or IRF3 5D (stimulus, 5 ng), different amounts of ORF3 expression constructs (l2.5 - l00 ng) and empty vector (pCG_HIV-l M NL4-3 *nef* stop ΔIRES-eGFP) to adjust total DNA amounts across all conditions to 200 ng/well. After 24 h, supernatants were harvested, and cells were lysed in l00µl lx Passive Lysis Buffer (Promega, Cat# El94A). *Gaussia* luciferase activity in the supernatants was measured by addition of Coelenterazine (PJK Biotech, Cat# l02l74). Firefly luciferase activity was measured in the cells using the Luciferase Assay System (Promega, Cat# El50l) according to the manufacturers instruction.

### LIPS assay

Luciferase fusion protein (l0^6^ RLU) in 50 µl Buffer A (50 mM Tris, 150 mM NaCl, 0.l% Triton X-100, pH 7.5) and 1 µl sample serum in 49 µl Buffer A was added to 1.5 ml tubes and incubated with shaking at 300 rpm for 1 h at room temperature. Pierce Protein A/G Magnetic beads were added to each condition as a 30% suspension in PBS for an additional hour and shaking at room temperature. Samples were placed on a magnetic rack, and supernatant was removed after 1-minute incubation. Magnetic beads were washed twice with 150 µl Buffer A followed by 2 washes with 150 µl PBS. Samples were transferred into a 96-well opaque nunc-plate (VWR), and 50 µl Coelenterazine (PJK Biotech, Cat# l02l74) was added to each condition. Samples were measured immediately on a TriStar2 S LB 942 Multimode Reader (Berthold Technologies) with an integration time of 0.1 seconds and a read height of 1 mm.

## QUANTIFICATION AND STATISTICAL ANALYSIS

Statistical analyses were performed using GraphPad PRISM 9.4.l. For statistical testing between two means P values were calculated using paired or unpaired Student’s t test. For comparison within one group, we used one-way analysis of variation (ANOVA) with Dunnett’s multiple comparison test and for comparison between two or more groups we used two-way ANOVA with Sidak’s multiple comparison test. Unless otherwise stated, data are shown as the mean of at least three independent experiments ± SD. Significant differences are indicated as: * p ≤ 0.05; ** p ≤ 0.01; *** p ≤ % 0.001 and **** p ≤ 0.0001. Statistical parameters are specified in the figure legends.

## ACKNOWLEDGMENTS

We thank Isabell HauBmann for excellent technical assistance and would like to thank the following persons for providing helpful reagents: Peter D Burbelo (LIPS assay plasmids), Konstantin MJ Sparrer (RIG-I, MDA5 and TBK1 expression plasmids, IRF luciferase reporter plasmid), Harmit Malik (MAVS expression plasmid), Rongtuan Lin (IRF3 expression plasmid), Bernd Baumann (NF-κB luciferase reporter plasmid, *Gaussia* luciferase control vector, IKKβ expression plasmid), Michael Gale Jr (IFN-β promoter reporter plasmid), Prof. Takasuke Fukuhara (CPER plasmids and cell lines) and Prof. Takashi Irie (pCAGGS Flag-IRF3 5d). SARS-CoV-2 mNeon Green was kindly provided by Prof. Michael Schindler. We would also like to thank Prof. Adolfo Garcia-Sastre and the CEIRR program (NIAD Centers of Excellence for Influenza Research and Response) for providing BiFC reporter plasmids. This work was funded by the Federal Ministry of Education and Research Germany (BMBF; grant ID: FKZ 0lKI20l35), the Canon Foundation Europe, the Heisenberg Program of the German Research Foundation (DFG; grant ID: SA 2676/3-1) and grants of the COVID-19 program of the Ministry of Science, Research and the Arts Baden-Württemberg (MWK; grants IDs: MWK K.N.K.C.0l4 (U2l) and MWK K.N.K.C.015 (U22)) to D.S. AE and AH were supported by BMBF SenseCoV2 0lKI20l72A; DFG Fokus COVID-19, EN 423/7-1; and Coronavirus research grants by the Bavarian State Ministry of Science and the Arts and Bavarian State Ministry of Health, Bay-VOC.

## AUTHOR CONTRIBUTIONS

M.M. and D.S. conceptualized the study and designed the experiments. M.M. performed most of the experiments. A.H. and A.E. generated and provided BAC clones for SARS-CoV-2 ΔORF6 YFP (ORF3c wild type and ΔORF3). S.F., M.M. and D.S. generated SARS-CoV-2 mutants via CPER that were characterized by D.S., S.F., M.M. and K.U.. C.K. and J.E.K. performed some of the luciferase reporter assays. M.S. sequenced the YFP reporter viruses. A.S. and J.I. performed *in silico* analyses to identify premature stop codons in ORF3c. M.M. and D.S. prepared the figures, and D.S. wrote the initial draft of the manuscript. All authors reviewed and edited the manuscript. A.E., K.S. and D.S. supervised the experiments, provided resources and acquired funding.

---

**Figure S1: Detection of SARS-CoV-2-specific antibodies via Luciferase Immunoprecipitation System (LIPS).**

**A** Principle of the LIPS assay: HEK293T cells are transfected with expression plasmids for a viral protein of interest fused to Renilla luciferase. Subsequently, transfected cells are lysed and incubated with serum samples and magnetic beads. Antibodies against viral proteins of interest will cross-link the luciferase-containing proteins with beads and allow magnet-assisted pull-down of both beads and luciferase activity.

**B.** LIPS-mediated quantification of antibodies against SARS-CoV-2 N (left panel) and ORF3c (right panel) in sera from SARS-CoV-2 naïve and convalescent sera (RLU, relative light units). Each dot represents one independent serum sample. Differences in antibody levels between SARS-CoV-2 naïve and convalescent sera was determined by unpaired student’s t-test with Welch’s correction; * p ≤ 0.05, ** p ≤ 0.01, *** p ≤ 0.001.

