## Supplementary figures and images for "ORF3c is expressed in SARS-CoV-2 infected cells and suppresses immune activation by inhibiting innate sensing"

### Supplemental Figure 1

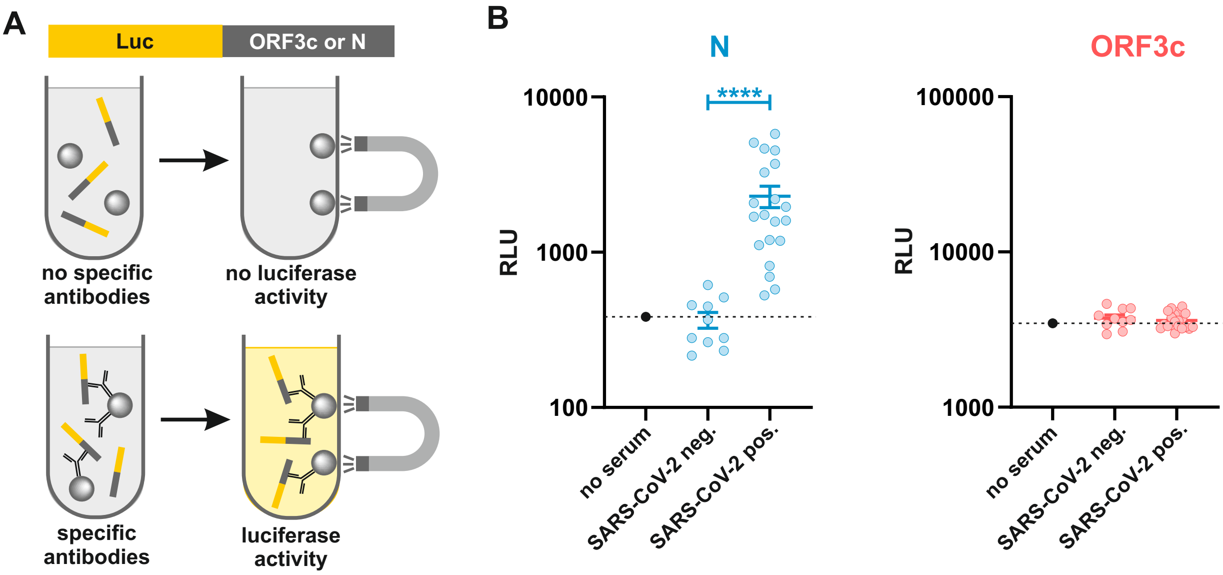
